# Environmental DNA sampling provides new management strategies for vernal pool branchiopods in California

**DOI:** 10.1101/2020.11.20.391052

**Authors:** Shannon Rose Kieran, Joshua Hull, Amanda Finger

**Affiliations:** Department of Animal Science, University of California, Davis, Davis, California, USA; Sacramento Fish and Wildlife Office, United States Fish and Wildlife Service, Sacramento, California, USA

## Abstract

California’s vernal pools are declining ecosystems that support valuable native plant and animal diversity. Vernal pool branchiopods are particularly at risk from vernal pool habitat loss and conservation efforts have targeted their long-term protection through the establishment of preserves and conservation banks. These conservation strategies require repeated, perpetual monitoring of preserved habitat, which is currently carried out through dip-net surveys and visual identification of specimens. Dip-netting may be destructive and frequently requires some sacrifice of protected species. Environmental DNA offers a new, modern method to monitor many protected freshwater organisms. We designed qPCR-based species-specific assays for four of California’s vernal pool branchiopods: The Vernal Pool Fairy Shrimp *Branchinecta lynchi* (BRLY), the Midvalley Fairy Shrimp *Branchinecta mesovallensis* (BRME), and the Conservancy Fairy Shrimp *Branchinecta conservatio* (BRCO), and the Vernal Pool Tadpole Shrimp *Lepidurus packardi* (LEPA). We tested these assays using eDNA sampling protocols alongside traditional dip-net surveys to assess their viability as an alternative method to monitor vernal pool branchiopods. Based on occupancy modeling, each of our assays achieved a 95% or higher detection rate when using optimized sampling protocols.

## INTRODUCTION

Vernal pool wetlands support high levels of biodiversity and provide a wide range of ecosystem services [1], but are facing decline. California’s vernal pool habitats support ecologically and phylogenetically distinct biota, including a diverse assemblage of endemic branchiopod crustaceans [2]. Since the mid-1800s, alteration of vernal pool wetlands and conversion to agricultural and urban landscapes are estimated to have contributed to the extinction of 15-33% of crustacean species in Central Valley vernal pools [3]. Of the remaining vernal pool crustacean species, six are listed as threatened or endangered [4].

A variety of conservation approaches have been implemented to help slow wetland loss and conserve remaining vernal pool biodiversity, with particular emphasis on conservation of vernal pool crustaceans. Conservation banks and habitat conservation plans are two important tools that establish permanent, managed habitat protections for vernal pools. These land ownership plans are designed to offset small-scale habitat loss from development and facilitate the protection of strategically-located, large, unfragmented habitats. A central part of these agreements are provisions for repeated, long-term monitoring of all listed species to help ensure their persistence within protected lands. Repeated monitoring ensures that the habitat bank perpetually supports the target, but comes at the cost of human impact on lands that are often pristine and delicate. Monitoring tools that minimize repeated impact would be useful for long-term monitoring of vernal pool conservation banks.

Currently, vernal pool branchiopods (Arthropoda: Crustacea: Branchiopoda) are primarily monitored during the wet season using dip-net surveys. Surveyors wade into vernal pools and move a fine mesh net through the water column to disrupt the substrate and capture benthic invertebrates. Once captured, adult vernal pool branchiopods are identified visually in-field or in the lab under a microscope. Drawbacks to this labor-intensive survey method include difficulty identifying morphologically-similar branchiopod species in-field, habitat disturbance, and death of specimens used for laboratory identification. Additionally, no detection rates have been established for dip-net surveys of adult vernal pool branchiopods, and juveniles in common target genera cannot be identified to species visually. Long-term monitoring plans would benefit from a survey method that was repeatable, with a known detection rate, that minimized the sacrifice of protected species while providing highly accurate results. Environmental DNA (eDNA) provides a viable alternative for sampling many habitats, including vernal pools, and monitoring species of conservation concern [16–22].

Here, we developed eDNA monitoring methods for three morphologically-similar vernal pool species of conservation concern: The Vernal Pool Fairy Shrimp *Branchinecta lynchi* (BRLY), the Midvalley Fairy Shrimp *Branchinecta mesovallensis* (BRME), and the Conservancy Fairy Shrimp *Branchinecta conservatio* (BRCO), as well as for the endangered Vernal Pool Tadpole Shrimp *Lepidurus packardi* (LEPA). We designed four species-specific qPCR assays to detect the presence of each of these four large branchiopods, which are commonly targeted and monitored in Central Valley dip-net surveys. We tested these protocols on tissue-derived DNA and using eDNA samples collected from existing conservation lands alongside dip-net surveys to determine their viability as an alternative monitoring method.

## RESULTS

### Laboratory Development and Validation

To develop qPCR assays able to identify our four species in the laboratory, we used Sanger sequences from whole specimens of each target species sourced from at least two sites in California. For each species we sequenced multiple genes to determine the best primer and probe sequences (“assay”) for each species (see *Materials and Methods*). Using individuals from multiple geographic sites allowed us to account for inter-population, intra-specific singlenucleotide polymorphisms (SNPs) which might impact the assay efficiency. We determined the best gene and assay independently for each species. We developed assays on the 12S Ribosomal RNA Gene for BRLY and BRME, the 16S Ribosomal RNA Gene for LEPA and Cytochrome Oxidase I for BRCO (see **Table 1** for the sequences of each assay). We tested the assays on tissue-derived target DNA and optimized our thermocycling protocols. Each assay amplifies tissue-derived target DNA within 20 cycles of qPCR, except for BRLY, which amplifies within 30 cycles of PCR. Our optimized reaction recipe used 1X Taqman Environmental Mastermix (ThermoFisher), 0.9 uM each forward and reverse primer, 0.15 uM probe, and 1X bovine serum albumin. The thermocycling protocol consisted of a 10 minute initial denaturing at 95 °C followed by 50 cycles of denaturing at 95 °C and a one minute elongation step that varied in temperature for each assay, see **Table 2** for full conditions.

**Table 1.**
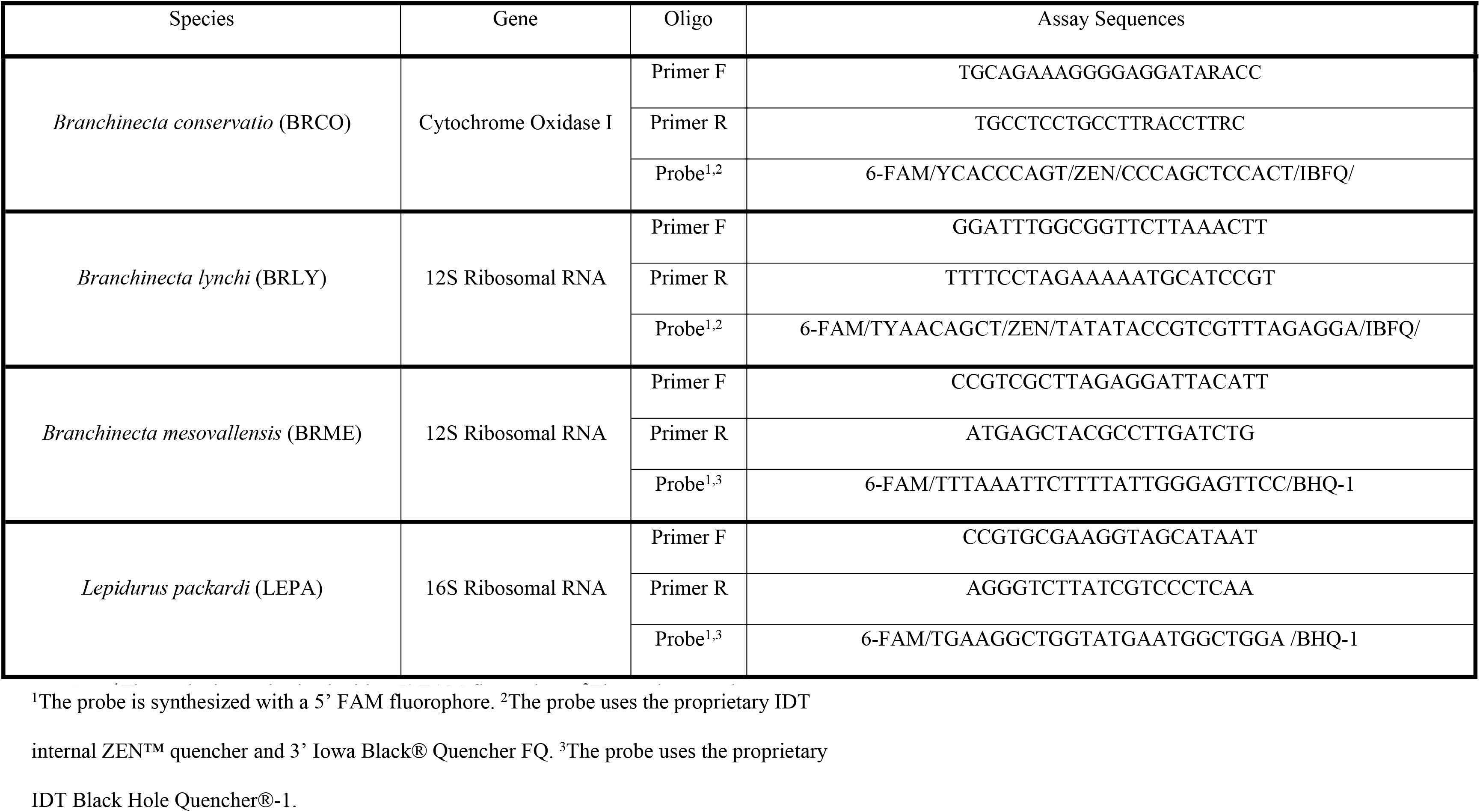
The complete DNA sequences of the primers and probes used in each species-specific assay.

**Table 2.**
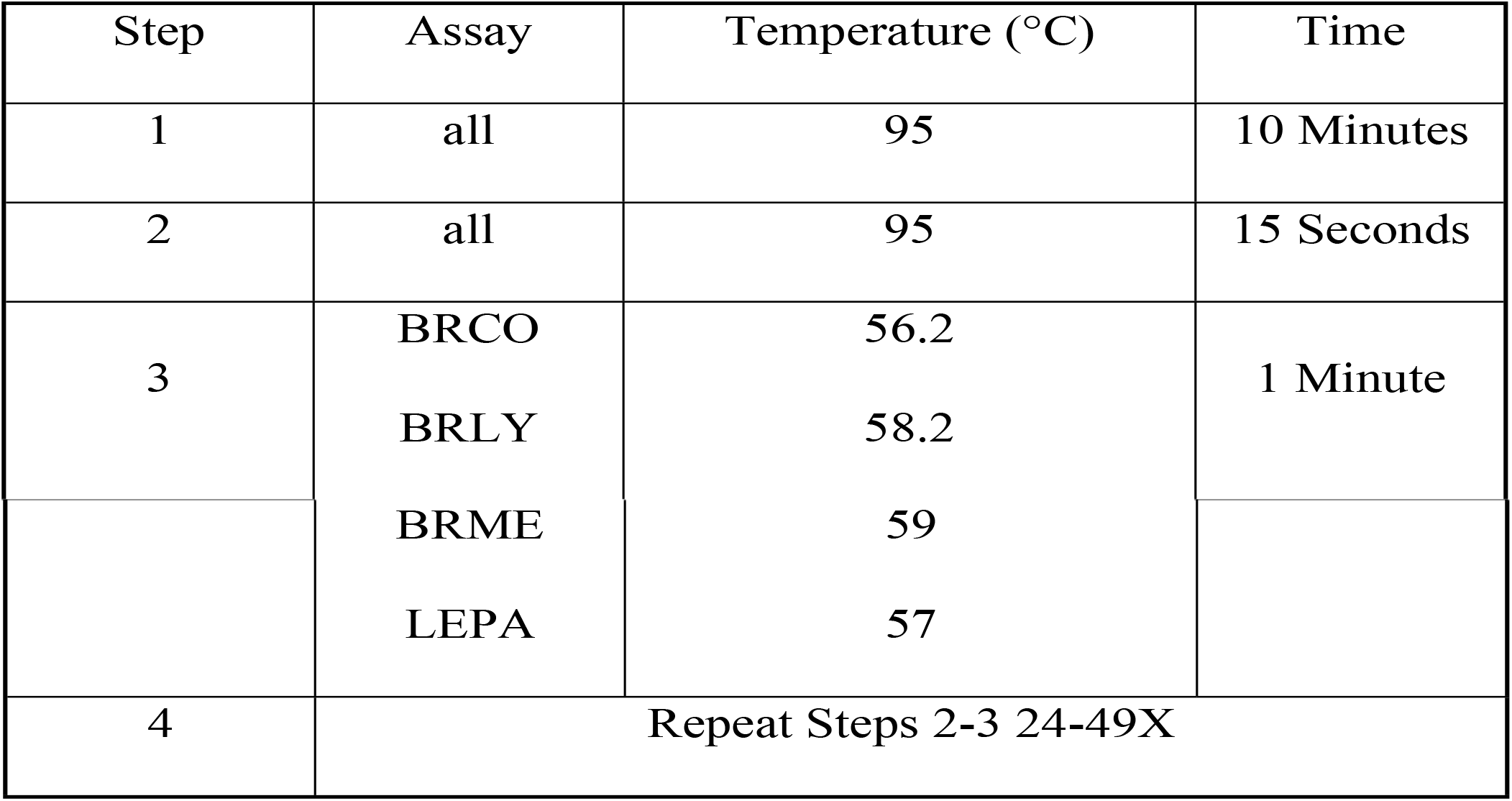
Thermocycling conditions for each of our assays.

To ensure that our assays were specific to our target species, we tested them using a panel of off-target, tissue-derived DNA that included every congeneric species known to occupy the same pools as any of our target species plus a frequent co-occurring branchiopod from the genus *Linderiella* (see **Table 3** for list, target species are in bold). There are no congeneric species whose ranges overlap with LEPA [5] so we tested the assay against the other branchiopod species in our off-target panel. We tested the assays using optimized thermocycling protocols for 50 qPCR cycles and found that no off-target species amplified within 30 cycles, while our targets all amplified before 20 cycles. This was true for all assays except BRLY, for which targets amplified within 30 cycles and no off-targets amplified before 42 cycles. We conclude that each assay is specific to its target when compared to a panel of co-occurring closely-related species run for 30 cycles (BRLY) or 20 cycles (BRME, BRCO and LEPA) of qPCR.

**Table 3.**
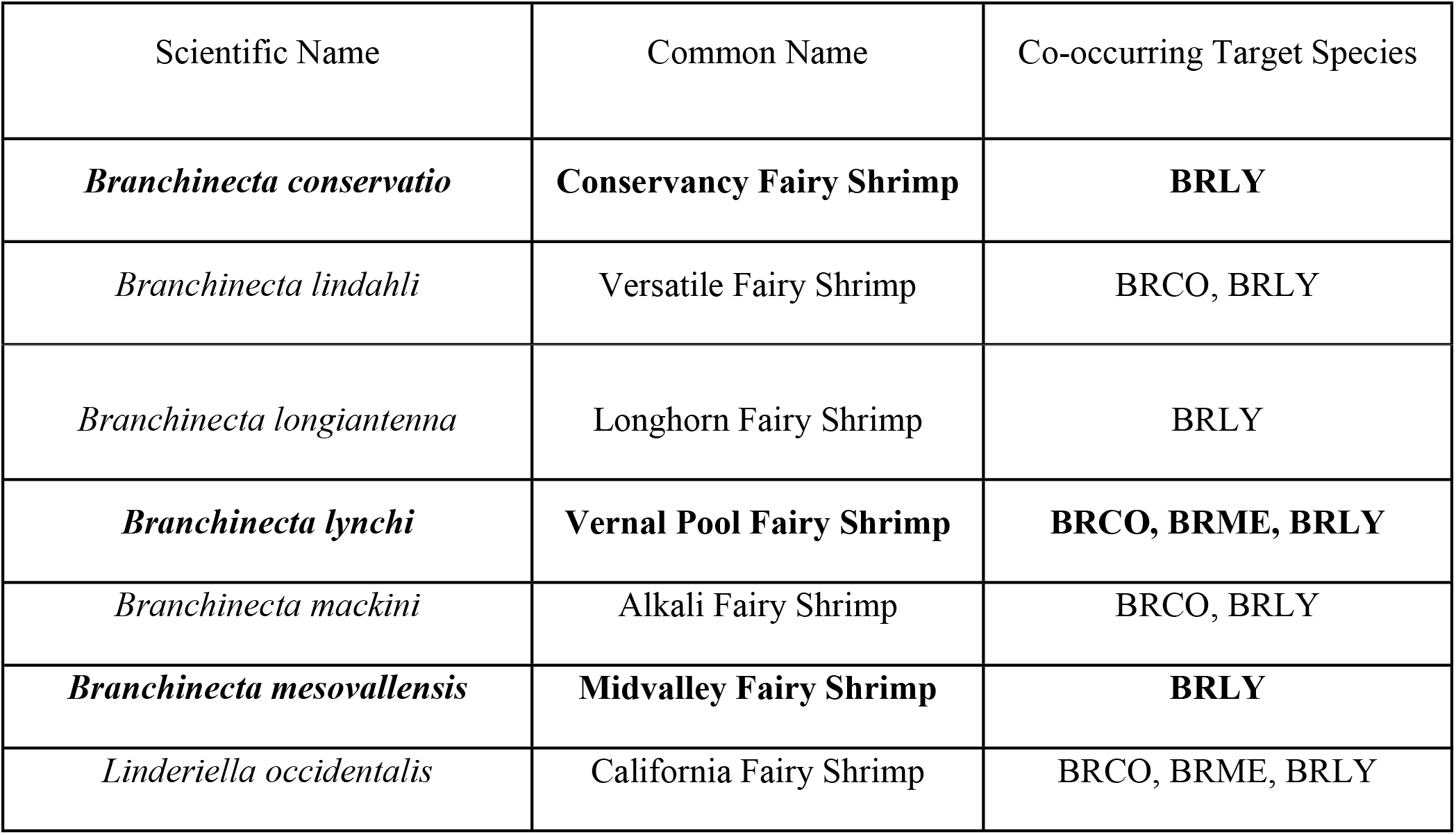
A list of off-target species used to test each assay, including the targets of our other assays tested to ensure no cross-amplification (in bold).

To determine the sensitivity of our assays, we used gBlock synthetic double-stranded oligos (IDT) matching the target amplicon of each assay at known concentrations. We produced a standard curve for each assay using five gBlock concentrations ranging from 1 copy/reaction to 100 copies/reaction with eight replicates at each concentration and eight no-template-controls per assay. We used the equation of each standard curve to calculate the limit of detection (LOD) and limit of quantitation (LOQ) for each assay. We found that the BRME assay was the most sensitive, with a calculated LOD of 5 copies/reaction and a calculated LOQ of 15 cycles/reaction, while our least sensitive assay, LEPA, had a calculated LOD of 12 copies/reaction and a calculated LOQ of 18 copies/reaction (see **Table 4**). A single animal cell may contain 5-10 copies of the mitochondrial genome [6], which suggests that our assays, under laboratory conditions, may be sensitive enough to detect the organism from the presence of a single cell. The concentration of target eDNA in the environment is often low [7], so a sensitive assay is important for use with environmental samples. We conclude that our assays are sensitive at low copy numbers in the laboratory and amplify when inputs are <20 copies/reaction of gBlock synthetic amplicons for all four target species.

**Table 4.** Limit of detection (LOD) and limit of quantitation (LOQ) for each assay as calculated from a standard curve using cycle quantification (Cq) mean and standard deviation.

| Assay | Cq Mean | Cq SD | Equation | LOQ | LOD | Efficiency | R<sup>2</sup> |
| --- | --- | --- | --- | --- | --- | --- | --- |
| **BRCO** | 35.23 | 0.444 | $y = -3.557x + 38.844$ | 18 | 10 | 91% | 0.919 |
| **BRLY** | 41.71 | 0.944 | $y = -3.61x + 44.343$ | 18 | 5 | 89.20% | 0.827 |
| **BRME** | 38.15 | 0.918 | $y = -3.5x + 40.442$ | 15 | 5 | 93% | 0.85 |
| **LEPA** | 36.69 | 0.424 | $y = -3.248x + 40.169$ | 18 | 12 | 103% | 0.722 |

### Field Validation

To evaluate the performance of our assays in the field, we collected eDNA samples from 89 vernal pools in the California Central Valley during a single wet season (see **Figure 1**). At each vernal pool, we collected three replicate 1 L water samples and then immediately dip-net surveyed the pool following U.S. Fish and Wildlife survey guidelines [8] (see *Materials and Methods*). Overall, we found high levels of agreement between our eDNA assays and dip-net survey results (see **Table 5** for summary). The BRLY assay was tested on 52 sampling events and agreed with positive dip-net results in 22 cases (85%). It agreed with negative dip-net results in 26 (100%) of cases. The BRME assay was tested on 56 sampling events and agreed with positive dip-net results in 13 (100%) cases and negative dip-net results in 43 (100%) of cases. The LEPA assay was tested on 45 sampling events. It agreed with positive dip-net results in 20 (91%) of cases. It agreed with negative dip-net results in 19 (86%) of cases. The BRCO assay was tested on 13 sampling events. It agreed with positive dip-net results in 5 (62.5%) cases. It was tested on 5 sampling events that had negative dip-net results and disagreed with negative dip-net results in 3 (60%) cases. However, all five of these sampling events came from pools with independently-confirmed presence of BRCO later that wet season. **S1 Table** contains a summary of sampling information. We conclude that all assays agree with positive dip-net results in the majority of cases.

**Figure 1.**
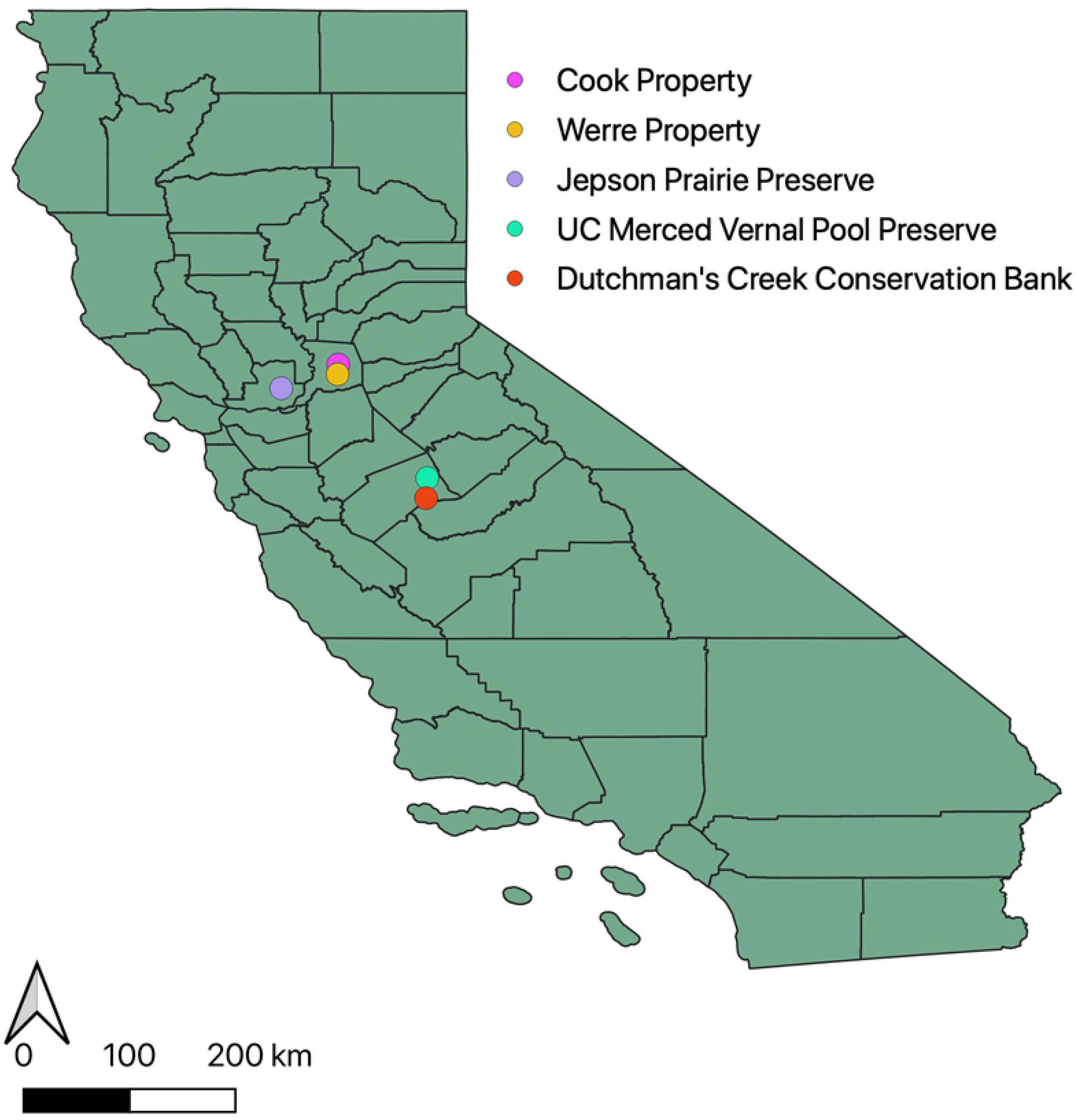
Map of properties sampled using side-by-side eDNA sampling and dip-net survey methods.

**Table 5.** Comparative results between eDNA (E) and dip-net (D) methods.

| Species | Total Sampling Events | E+/D+ | E-/D- | E-/D+ | E+/D- |
| --- | --- | --- | --- | --- | --- |
| BRCO | 13 | 5/8 (62.5%) | 2/5* (40%) | 3/8 (37.5%) | 3/5* (60%) |
| BRLY | 52 | 22/26 (84.6%) | 26/26 (100%) | 4/26 (15.4%) | 0/26 (0%) |
| BRME | 56 | 13/13 (100%) | 43/43 (100%) | 0/13 (0%) | 0/43 (0%) |
| LEPA | 44 | 20/22 (90.9%) | 19/22 (86%) | 2/22 (9.1%) | 3/22 (13.6%) |

\*These pools were independently determined to contain BRCO, despite negative dip-net results.

To estimate the detection rates of each assay, we carried out occupancy modelling for each target species using the side-by-side dip-net and eDNA survey results. We used the *R* statistical software and the R package *Unmarked* [9] to produce single-season occupancy models for each species. We provided *Unmarked* with four variables that might affect detection rates to use as covariates of detection: water volume filtered (per replicate), average water volume filtered (across replicates), pool area, and filtration protocol (in-field or in-lab). We produced graphs representing the probability of detecting a target in at least one replicate water sample using the R package ggplot2 [10]. For the BRLY and BRME assays, we found the base model (containing no covariates of detection) fit best, suggesting that these assays are robust to sampling protocols. Our model suggests that when the target species is present in a vernal pool, a single water replicate will detect BRLY 75.64% of the time (95% CI: 65.11-85.17%) and detect BRME 97.44% of the time (95% CI: 96.43-100%), see **Figure 2.** We conclude that these two assays effectively detect their target species at least 95% of the time with as few as one water replicate (BRME) or three water replicates (BRLY).

**Figure 2.**
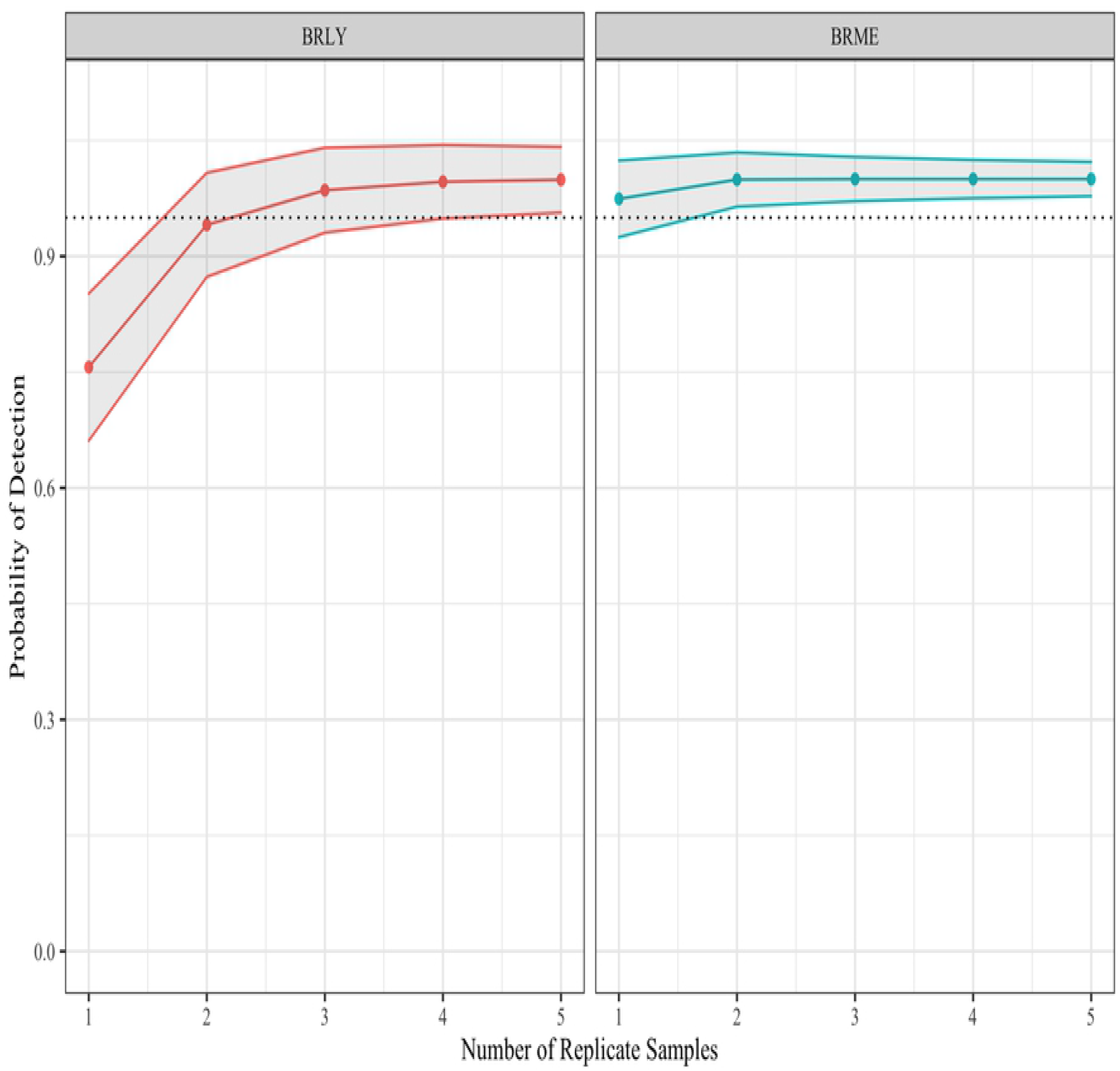
Probability of detection for BRLY and BRME using eDNA sampling with varying numbers of replicate water samples. Dotted line represents 95% detection probability.

In contrast to BRME and BRLY, our LEPA assay was sensitive to filtration protocols. Our comparative model fitting suggests that for LEPA, vernal pool area and filtration protocol (laboratory vs. field) were important covariates of detection. Additionally, the detection rate increased with increasing pool area. Laboratory filtration was consistently superior to field filtration. **Figure 3a-e** describes this. Each graph **a, b, c, d**, and **e** represents increasing vernal pool area size after normalization: a pool of area −1 has an area one standard deviation lower than the mean, a pool of area 0 is an average pool and a pool of area 1 is one standard deviation larger than the mean. Pools varied in size from 10 m^2^ to more than 10,000 m^2^. Although LEPA is sensitive to both filtration protocol and pool area, the assay can detect the target species in any pool size with 98.67% accuracy if at least five water samples are collected (95% CI: 88.42-100%) and filtered in the laboratory. Increasing replicates was not enough to reach a sufficient detection probability in very small pools using field filtration.

**Figure 3.**
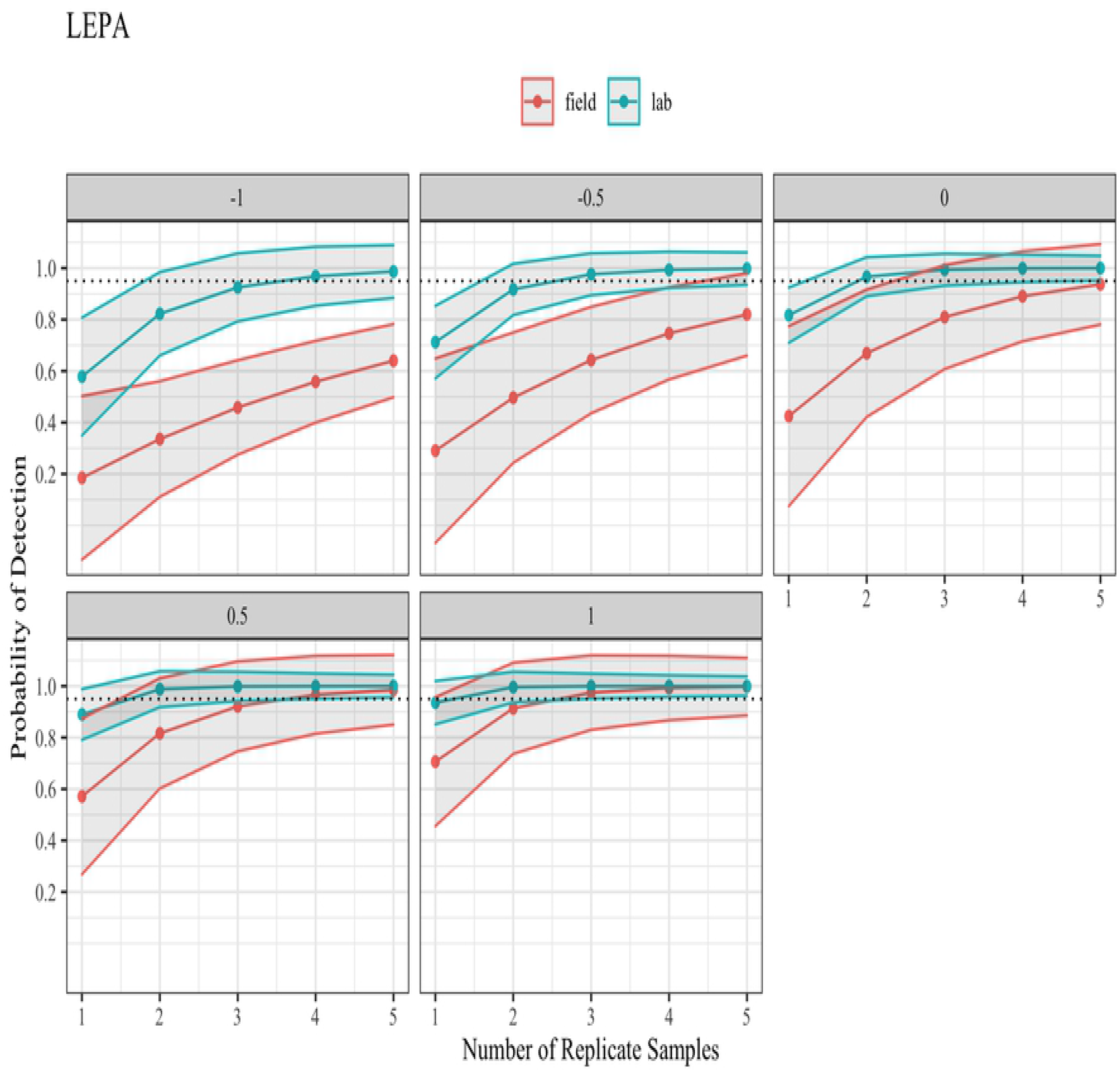
Probability of detecting LEPA using eDNA sampling in pools of varying size, per water replicate sampled. Dotted line represents 95% detection probability.

Because BRCO is a relatively rare species, we added an extra year of sampling in 2018 at three vernal pools to produce additional data. We used a larger-pore filter which allowed us to filter an increased volume of water, as pools at this site have high turbidity (see *Materials and Methods*). Our comparative model fitting suggests that for BRCO, average volume of water filtered and filtration protocol were the important covariates of detection. Our models did not suggest that the change in filter type or pore size affected detection rates. The BRCO assay detection rate was positively correlated with average water volume filtered, with very low detection rates for samples of 50 mL or less and much higher success rates with 1 L samples. Surprisingly, the filtration protocol seems to strongly favors in-field filtration. If samples are lab-filtered and water samples are <1 L, there is no number of replicate water samples that ensures a >95% detection rate. **Figure 4a-d** compares the assay detection rates at (**a**) 50 mL, (**b**) 100 mL, (**c**) 500 mL and (**d**) 1 L average water volume filtered per sample. The assay clearly performs better when filtration is immediate, although the difference is minimal at high (>1 L) volumes of water. To achieve >95% detection rate, five replicate field-filtered water samples are required in very turbid pools where water volume per sample is expected to be ≤50mL, but in clear pools where higher water volume per sample is expected, >99% detection can be achieved with only one field-filtered water replicate (95% CI: 99.70-100%) or three lab-filtered water replicates (95% CI: 89.89 −100%).

**Figure 4.**
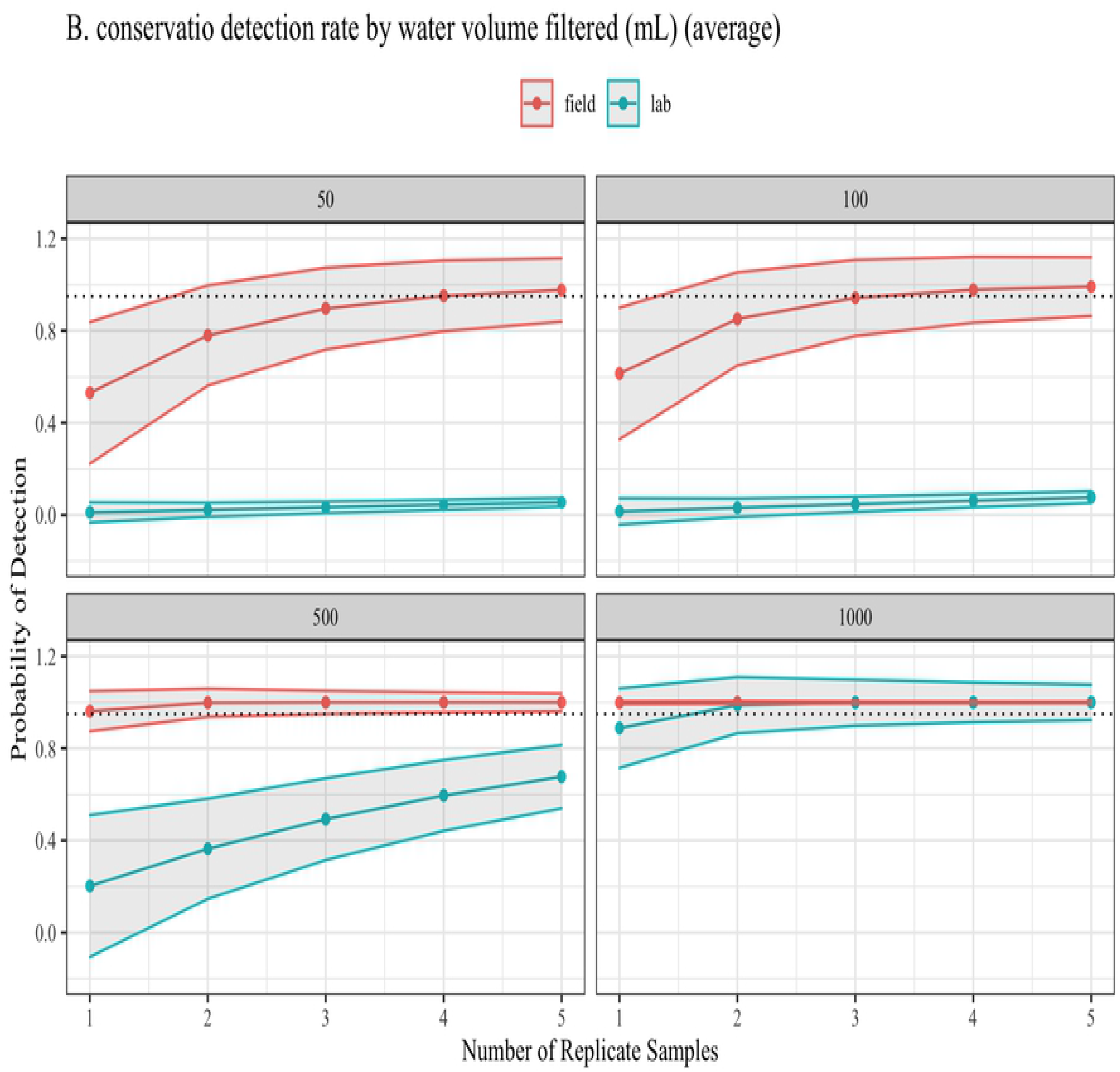
Probability of detecting BRCO using eDNA sampling using varying water volumes and water replicate samples. Volumes are measured in mL. Dotted line represents 95% detection probability.

To determine the false positive rate, we compared our dip-net and eDNA survey results for each species and investigated cases where the dip-nets did not detect our target, but the eDNA assay returned a positive response. We found few false positives in our data. There were no instances of positive eDNA survey results when dip-net results were negative in our BRLY or BRME datasets. There were three instances where the LEPA assay detected the presence of the target species but dip-nets did not find them. We examined these three sampling events in depth and found that in all three of them, the species was found in that pool either later in the same year or in other years. This leads us to believe that the assay is very sensitive and may out-perform dip-net surveys, but would benefit from expanded testing at sites known to be historically negative for LEPA. The BRCO assay had three sampling events where the eDNA assay detected BRCO but the dip-nets did not. All three of these sampling events were from three pools sampled in early 2018. The species was independently confirmed to be present in all three of these pools later in the year. These pools are large and turbid, and these sampling events represent cases where the eDNA assay appears to be more sensitive than dip-net methods. We conclude that the assays did not produce false positive results.

## DISCUSSION

We designed qPCR assays that reliably assess the presence of four California vernal pool branchiopod species using either tissue or environmental DNA samples. We targeted *Branchinecta lynchi* (BRLY), *Branchinecta conservatio* (BRCO), *Branchinecta mesovallensis* (BRME) and *Lepidurus packardi* (LEPA) because they are common targets of dip-net monitoring surveys and are protected species, or, in the case of BRME, coexist with and closely resemble protected species. Despite their protected status, few resources have been developed to support the long-term conservation of these species, and this is the first time that specific molecular markers have been published for any of these organisms. We designed these assays to maximize their utility and provide diverse applications in the field, but particularly aimed to support the long-term survey efforts required by conservation plans established for conservation banks and protected lands.

Our assays can be used with tissue-derived DNA for species-level identification of specimens collected by dip-net surveyors. Though visual identification error rates have not been established, *Branchinecta* species are phenotypically plastic and morphologically similar. Accurate visual identification can be difficult and likely varies with surveyor experience. DNA-based identification offers an objective method of differentiation that may improve survey data quality. Additionally, immature *Branchinecta spp*. are not visually distinguishable even under a microscope, which restricts dip-net surveys to the brief period when adult shrimp are occupying the pools. By applying these assays to preserved immature fairy shrimp specimens, it may be possible to expand dip-net surveys earlier in the field season. Alternatively, eDNA applications of these assays may reduce the need for dip-net surveys altogether.

A method that is non-lethal, repeatable and objective may be of particular value for the long-term monitoring of vernal pool habitats. Our method can be used without causing injury or mortality to protected species, and minimizes the destruction of the sensitive, high quality vernal habitat typically preserved by land-based conservation strategies. This study also provides the only quantitative detection rate for any vernal pool branchiopod survey method. Each of our assays achieved greater than a 95% detection rate using optimized eDNA sampling protocols. This suggests that these assays can be used as part of a monitoring plan to reliably and repeatably detect our targets.

These assays contribute to a growing library of resources to support the long-term maintenance of protected vernal pool species. Recent work has used eDNA metabarcoding methods to investigate branchiopod communities in Southern California [11] and to survey protected amphibians that breed in vernal pools [12]. These projects reflect a demand for modern monitoring methods. Future improvements to eDNA assay and survey methodology may be able to further improve management of California’s vernal pools using emerging eDNA technologies such as SHERLOCK [13], while population and landscape genetic data may provide managers with new genetic management tools for these vulnerable populations.

Vernal pool habitats continue to decline and the USFWS Recovery Plan calls for long-term, repeated monitoring of currently-conserved habitat alongside the perpetual conservation of at least 80% of suitable habitat for target species [14]. Repeatable, objective methods could vastly increase the quality of species presence data from protected vernal pool habitats, which in turn will facilitate effective long-term preservation of these ecosystems. This work provides new tools for managers in California’s Central Valley to survey and track populations of protected branchiopods using methods with known detection rates and minimal sacrifice of specimens.

## MATERIALS AND METHODS

### Assay Design and Validation

We used both archived and fresh specimens when developing our assays. Archived tissue was sourced from the Bohart Museum of Entomology in Davis, CA and from the USFWS Sacramento Regional Office in Sacramento, CA. We extracted DNA from ethanol-stored specimens using the DNEasy Blood and Tissue Kit (Qiagen) following a modified protocol from the manufacturer. Our first modification was to rinse each specimen in DI water and air-dry it for 10 minutes on a Kimwipe to remove excess ethanol. Following the rinse and dry step, we removed the intestines and mature egg sacs with scalpels and forceps. All equipment was sterilized using bleach and DI water between specimens. Our second modification was to incubate specimens on a rotisserie for 12 hours or overnight after the addition of Qiagen Buffer ATL and proteinase K. Extraction continued as recommended by the manufacturer, using the optional second elution step for a total of two 60 μL elutions.

We Sanger sequenced our extracted DNA using preciously-published primers provided in **S2 Table** to assess multiple genes [15–18]. We aligned the sequences produced by each set of primers using MEGA7 [19] to produce a consensus sequence which we input into Primer3Plus [20] and PrimerQuest (IDT) which produced candidate assays. We then visually inspected the candidate assays produced by each program against our aligned sequences to look for inter-specific and intra-specific SNPs. We selected candidate assays that maximized inter-specific differences without introducing intra-specific SNPs into our assay. We analyzed each candidate assay with IDT’s Oligo Analyzer software to account for the change in melting temperature expected after the addition of the fluorophore and quencher molecules, as well as identify potential secondary structures which could hamper assay efficiency. To optimize the qPCR protocols for our new assays, we used our previously-extracted tissue-derived target DNA and ran a gradient PCR to determine the optimum annealing temperature for each assay.

To calculate the Limit of Detection (LOD) and Limit of Quantitation (LOQ), we used gBlock double-stranded gene fragments (IDT) that matched the target amplicon of each of our target species as the input for each assay. We used gBlocks because their precise molecular weight allows us to more accurately calculate copies/reaction. An assay that uses a gBlock standard as an input may be more sensitive than one that uses tissue-derived DNA, because the gBlock standard is a short fragment that amplifies with maximum efficiency. Thus, the results of our LOD and LOQ calculations represent the assay’s performance under “ideal” conditions. In real life conditions, large molecules of DNA may be less efficient, and environmental samples introduce potential PCR inhibitors which may also lower efficiency, but accurate LOD and LOW must be derived from known input concentrations, which are not possible to determine from eDNA, as DNA concentration in an eDNA sample is not an accurate measure of target species DNA concentration. We produced a standard curve with concentrations that ranged from 100 copies/reaction to 1 copy/reaction. The LOD is defined as the lowest concentration of analyte that can be reliably differentiated from a blank at 95% confidence, while the LOQ is defined as the lowest concentration that can be accurately quantified. We used the formulas for LOD and LOQ developed by the USGS Ohio Water Microbiology Laboratory [21].

### Field sampling

We sampled 89 vernal pools for branchiopod presence during the 2017 wet season and an additional three vernal pools during the 2018 wet season. To collect environmental samples from a vernal pool, we collected water into a sterile 1L Nalgene bottle. We took three replicate water samples at each vernal pool and each water sample was collected from a different area of the pool. After eDNA sampling we immediately dip-net surveyed the pool and collected pool area measurements by walking the perimeter of the pool using a GPS device with an area tracking feature (Garmin). We followed the USFWS guidelines for large branchiopod surveys and collected voucher specimens in 95% ethanol under USFWS Recovery permit FWSSFWO-16. When possible, voucher specimens were independently identified to species.

### Filtration

Filtration of eDNA samples was carried out by pouring the water from the screw-top Nalgene over a single-use cellulose nitrate filter (0.22 uM, 47 mm diameter, Steriltech) housed in a single-use plastic filter housing. The filter and housing were attached to a 1 L vacuum flask which was attached via rubber tubing to a peristaltic pump. Water was continually poured into the filter housing. Water volume filtered was not varied intentionally, but filtration was ended when the filter clogged or 500 mL was filtered (up to 1000 mL in 2018).

To determine if immediate, in-field filtration of our water samples increased our detection rates, we tested both in-field and in-laboratory filtration protocols. In the field, filtration took place immediately after environmental sample collection at the side of the vernal pool, before dip-net surveys were carried out. To detect any possible contamination when filtering in the field, a “field negative” control consisting of 500 mL sterile nanopure water was carried into the field in a closed screw-top Nalgene and filtered immediately following the filtration of the three replicate water samples. The negative control was included in extraction and qPCR analysis identically. Sampling events where the negative control amplified for any species were thrown out. Laboratory-based filtration took place immediately after returning from the field. After environmental sample collection, the screw-top Nalgenes containing our water samples were placed in individual sealed 1-gallon zip-top bags. Bags were placed on ice in a cooler and returned to the filtration laboratory within 4 hours, at which time they were immediately placed into a 4 °C refrigerator and filtered within 12 hours following the same protocol as above. An “equipment negative” identical to a field negative was included for lab filtration.

In 2018, we tested a larger-pore glass fiber filter on three pools at Jepson Prairie Preserve to increase the number of BRCO samples in our data. We visited three vernal pools three times each for nine total samples (27 replicate water samples) and dip-net surveyed each pool after each visit. We performed all filtration in the laboratory and filtered up to 1 L of water, and no voucher specimens were collected. Filter storage and processing occurred exactly as in 2017.

### Environmental DNA Extraction

To extract DNA from our filtered eDNA samples, we used the Qiagen DNEasy Blood and Tissue Kit using the following modifications. 1) Because filtered samples were stored in 180 uL Buffer ATL, Proteinase K was added directly to this tube. 2) Samples were incubated on a rotisserie for 12 hours or overnight. 3) When samples were transferred to columns, care was taken to avoid transferring any remaining filter, as filters did not disintegrate fully. 4) Instead of a single elution, two elutions of 60 mL each were performed, and 5) a 15-minute incubation of the eluent at 55 °C (rather than one minute at room temperature) was performed during each elution step. Finally, to prevent PCR inhibition, after extraction, all samples were proactively treated for inhibitor removal using the Zymo One-Step PCR Inhibitor Removal Kit. All consumables including microcentrifuge tubes, forceps, pipette tips and qPCR plates and water were sterilized before use in a UV crosslinker for 10 minutes.

### qPCR Analysis

Field replicate samples were assayed separately. Each sample and replicate was run in quadruplicate and each qPCR plate included two low-concentration gBlock positive controls and a no-template control, each also run in quadruplicate. Each qPCR plate tested for only one target species using the optimized thermocycling conditions for 50 cycles. If one qPCR replicate of the four amplified, the sample was re-run. If two or more amplified, the sample was considered positive. If zero amplified, the sample was considered negative. If one replicate of four amplified after being re-run a second time, it was considered positive.

### Contamination Controls

We attempted to control for contamination at every step in the process. During environmental sample collection, collectors wore single-use nitrile gloves when handling the bottles and the filtration mechanisms, which were changed between pools. Collectors wore sterile, single-use boot covers when they had to wade into the pools. The 1 L Nalgene collection bottles were sterilized between uses by soaking in 20% bleach for 30 minutes, followed by triple rinsing and 15 minutes sterilization under a UV hood. All reusable filtration equipment was sterilized between uses following the same method. Boots and nets were cleaned between sites following USFWS decontamination procedures. We maximized our single-use equipment, choosing single-use filter housings, filter storage tubes, zip-top bags and forceps. Filtration was carried out in a designated laboratory space that contained no PCR product or extracted DNA, but was not our clean laboratory (as field gear and filtration supplies would contaminate a clean laboratory).

All post-filtration steps were carried out in a clean laboratory, where no PCR product or tissue-derived DNA was permitted. Personnel were not permitted to move between the clean laboratory and any laboratory containing crustacean tissue, tissue-derived DNA or PCR product without first changing clothes and shoes. No reagents, equipment or consumables were permitted into the clean laboratory from any laboratory containing crustacean tissue, tissue-derived DNA or PCR product. Within the clean laboratory, all work took place in a UV hood.

During DNA extraction an “extraction blank”, a tube of containing only nanopure water, was carried through the entire extraction process alongside our samples. These blanks were tested with all four assays to ensure the extraction process did not introduce contaminants. We used filter-tip pipette tips for all work. All consumables (microcentrifuge tubes, falcon tubes, qPCR plates, etc.) were UV sterilized for 10 minutes in a UV crosslinker before use, along with all water used in DNA extraction or qPCR. Reusable equipment, such as tube racks, forceps, and beakers, were bleach sterilized by soaking for 20 minutes in 20% bleach and triple rinsing in DI water, followed by 10 minutes in the UV crosslinker. After setting up qPCR reactions, plates were sealed and removed to another, non-clean space still free of tissue-derived or amplified crustacean DNA, where qPCR was carried out on a Bio-Rad CFX96.

### Modeling

We developed our models using the R package *Unmarked*. We treated each sampling event (consisting of three replicate water samples and a dip-net survey) as an independent event. We produced models for every combination of our four covariates of detection (water volume filtered per replicate, average water volume filtered, filtration location and pool size) and used AIC-based model selection to select the best model for each species. We used the MacKenzie and Bailey goodness-of-fit test [22] on each selected model to determine appropriate fit. We treated sampling events with positive dip-net results for a target species as a known occupied site for that species. We found that without this additional information, the models greatly exaggerated our detection rates, likely due to the high level of concordance between our eDNA field replicates. To visualize the model results, we used *Unmarked’*s “predict” function to estimate detection rates at specific relevant parameters.

## ACKNOWLEDGMENTS

The findings and conclusions in this article are those of the author(s) and do not necessarily represent the views of the U.S. Fish and Wildlife Service. The authors would like to acknowledge the invaluable assistance of Alisha Goodbla, Alyssa Benjamin, Amanda Coen, Henry Hwang, Kaitlyn McGee, Pauline Tran, Jason Peters and Daniel Prince for their help collecting samples. We would like to acknowledge Carol Witham for her work identifying voucher specimens. We would also like to thank our partners at our field sites: Virginia Boucher with The University of California Natural Reserve System, Tara Collins with Westervelt Ecological Services, Carly Rich with ECORP Consulting, and Lucie Adams with the Sacramento Valley Conservancy. We would like to especially thank Monique Kolster with the University of California, Merced, for her tireless advocacy work on behalf of vernal pool research. This project was funded primarily through U.S. Bureau of Reclamation CESU R15AC00040, with additional funding through the University of California Natural Reserve System.

## SUPPLEMENTAL TABLE CAPTIONS

**S1 Table. Detailed eDNA sample collection data. Columns include a unique collection ID, vernal pool ID, the property sampled, the filter type, filtration protocol, sampling date, pool area in square meters, water volume sampled (per water replicate) in milliliters, and the latitude and longitude of the pool in decimal degrees.**

**S2 Table. Primers used to Sanger sequence tissue samples for assay development.**

## REFERENCES

1. Duffy WG, Kahara SN. Wetland ecosystem services in California’s Central Valley and implications for the Wetland Reserve Program. Ecol Appl. 2011;21: 18–30. doi:10.1890/09-1338.1

2. King JL, Simovich MA, Brusca RC. Species richness, endemism and ecology of crustacean assemblages in northern California vernal pools. Hydrobiologia. 1996;328: 85–116. doi:10.1007/BF00018707

3. King J. Loss of diversity as a consequence of habitat destruction in California vernal pools. Ecology, Conservation, and Management of Vernal Pool Ecosystems. 1998. pp. 119–123. Available from: http://www.vernalpools.org/proceedings/king.pdf

4. California Endangered Species Act. United States; 1973. Available from: https://leginfo.legislature.ca.gov/faces/codes_displayText.xhtml?lawCode=FGC&division=3.&title=&part=&chapter=1.5.&article=1.

5. Rogers DC. Revision of the nearctic Lepidurus (Notostraca). J Crustac Biol. 2001;21: 991–1006. doi:10.1651/0278-0372(2001)021[0991:rotnln]2.0.co;2

6. Moraes CT. What regulates mitochondrial DNA copy number in animal cells? Trends in Genetics. 2001. doi:10.1016/S0168-9525(01)02238-7

7. Song J. Making Sense of the Noise: Statistical Analysis of Environmental DNA Sampling for Invasive Asian Carp Monitoring Near the Great Lakes. Carnegie Mellon Univ. 2017. Available from: http://repository.cmu.edu/dissertations/901

8. United States Fish and Wildlife Service. Survey Guidelines for Listed Large Branchiopods. 2015. Available from: https://www.fws.gov/sacramento/es/Survey-Protocols-Guidelines/Documents/VernalPoolBranchiopodSurveyGuidelines_20150531.pdf

9. Fiske I, Chandler R. Overview of Unmarked: An R Package for the Analysis of Data from Unmarked Animals. R. 2015; 1–5. doi:10.1002/wics.10

10. Wickham H. ggplot2. ggplot2. 2009. doi:10.1007/978-0-387-98141-3

11. Gold Z, Wall AR, Curd EE, Kelly RP, Pentcheff ND, Ripma L, et al. eDNA metabarcoding bioassessment of endangered fairy shrimp (Branchinecta spp.). Conserv Genet Resour. 2020. doi:10.1007/s12686-020-01161-9

12. Kieran S, Hull JM, Finger AJ. Using environmental DNA to monitor the spatial distribution of the California Tiger Salamander. J Fish Wildl Manag. 2020; 11: 1. doi:https://doi.org/10.3996/052019-JFWM-041

13. Williams MA, O’Grady J, Ball B, Carlsson J, de Eyto E, McGinnity P, et al. The application of CRISPR-Cas for single species identification from environmental DNA. Mol Ecol Resour. 2019. doi:10.1111/1755-0998.13045

14. United States Fish and Wildlife Service. Recovery Plan for Vernal Pool Ecosystems of California and Southern Oregon. 2005. Available from: https://www.fws.gov/sacramento/es/Recovery-Planning/Vernal-Pool/

15. Machida RJ, Kweskin M, Knowlton N. PCR primers for metazoan mitochondrial 12S ribosomal DNA sequences. PLoS One. 2012. doi:10.1371/journal.pone.0035887

16. Folmer O, Black M, Hoeh W, Lutz R, Vrijenhoek R. DNA primers for amplification of mitochondrial cytochrome c oxidase subunit I from diverse metazoan invertebrates. Mol Mar Biol Biotechnol. 1994;3: 294–299. doi:10.1371/journal.pone.0013102

17. Deiner K, Hull JM, May B. Range-wide phylogeographic structure of the vernal pool fairy shrimp (Branchinecta lynchi). PLoS One. 2017; 1–20.

18. Apakupakul K, Siddall ME, Burreson EM. Higher Level Relationships of Leeches (Annelida: Clitellata: Euhirudinea) Based on Morphology and Gene Sequences. Mol Phylogenet Evol. 1999. doi:10.1006/mpev.1999.0639

19. Kumar S, Stecher G, Tamura K. MEGA7: Molecular Evolutionary Genetics Analysis Version 7.0 for Bigger Datasets. Mol Biol Evol. 2016;33: 1870–1874. doi:10.1093/molbev/msw054

20. Untergasser A, Nijveen H, Rao X, Bisseling T, Geurts R, Leunissen JAM. Primer3Plus, an enhanced web interface to Primer3. Nucleic Acids Res. 2007;35: W71–W74. doi:10.1093/nar/gkm306

21. Francy DS, Bushon RN, Cicale JR, Brady AMG, Kephart CM, Stelzer EA, et al. Quality Assurance/Quality Control Manual: Ohio Water Microbiology Laboratory. Columbus, OH; 2017. Available from: https://oh.water.usgs.gov/OWML/micro_qaqc.htm

22. MacKenzie DI, Bailey LL. Assessing the fit of site-occupancy models. J Agric Biol Environ Stat. 2004. doi:10.1198/108571104X3361

